# Capsule-mediated O-antigen masking protects hypervirulent *Klebsiella pneumoniae* from neutrophil killing

**DOI:** 10.1101/2025.02.26.640295

**Authors:** Karin Santoni, Joshua L.C Wong, Renaud Poincloux, Etienne Meunier, Gad Frankel

## Abstract

*Klebsiella pneumoniae* is listed as a critical priority pathogen by the WHO due to prevalent extended-spectrum β-lactamase and carbapenem resistance and high mortality rates Hypervirulent *K. pneumoniae* (hvKp) causes life-threatening infections in immunocompetent individuals, while classical *K. pneumoniae* (cKp) causes nosocomial infections in immunosuppressed patients. The convergence of hypervirulence with antibiotic resistance is now a major global health threat. Using a murine pulmonary infection model, we show that hvKp resisted clearance despite significant neutrophil recruitment. Using clinical hvKp and cKp isolates, bone marrow derived neutrophils, cell biology, pharmacology and imaging (both light and electron microscopy) in ex vivo assays revealed that the hvKp hypermucoviscous capsule prevents bacterial recognition and neutrophil activation, blocking ROS production, degranulation, and neutrophil extracellular traps (NETs) formation. In the absence of capsule (Δ*wcaJ*) or hypermucoid capsule determinants (Δ*rmpADC*), hvKp activates neutrophils, leading to degranulation, NET formation and bacterial killing. While superimposing the absence of O-antigen (Δ*rfb*) on the Δ*wc*aJ abrogates degranulation, neutrophils still elaborate NETs that eliminate the Δ*wcaJ*/Δ*rfb* mutant at high efficiency. This suggests that O-antigen is a double-edged sword, functioning as a PAMP triggering neutrophil degranulation yet assisting the pathogen to evade NET-mediated destruction. These findings shed light on the mechanism underpinning immune evasion by the hypervirulent capsule in hvKp.

## Introduction

*Klebsiella pneumoniae* (Kp) is an extracellular Gram-negative pathogen, belonging to the Enterobacteriaceae family^1^. Kp is listed as a critical priority pathogen by the WHO due to prevalent extended-spectrum β-lactamase and carbapenem resistance and high mortality rates^2,3^. It is responsible for about one-third of all Gram-negative healthcare-associated infections including pneumonia, surgical wound, urinary tract and bloodstream infections. These infections can result in sepsis and septic shock, with a high attributable mortality approaching 30%^3^. Worldwide, Kp is responsible for 30 million infections and more than 500,000 deaths annually^2,3^.

Kp is broadly classified into classical opportunistic multi-drug resistant Kp (cKp) - causing nosocomial infections in immunosuppressed patients - and hypervirulent Kp (hvKp), causing community-acquired infections in immunocompetent hosts^1,4^. The convergence of hypervirulence with antibiotic resistance is now considered a major global public health concern^1,5^. HvKp strains typically exhibit a hypermucoviscous (HMV) phenotype and thicker capsular polysaccharide (CPS) compared to cKp^4,6,7^. The hypercapsulation and hypermucoviscosity phenotypes are driven by genes encoded in the *rmpADC* locus (regulator of the mucoid phenotype)^8,9^. The hypervirulent capsule is a known virulence factor enabling hvKp to evade innate immunity^1,7,10^.

The first responder innate immune cells are neutrophils, which represent about 70% of all peripheral leukocytes. Neutrophils are highly effective phagocytes, capable of eliminating up to two times more pathogens than other phagocytic cells^11,12^. They capture pathogens in a phagosome, which then fuses with granules filled with antimicrobial substances^13^. Phagocytosis is also accompanied by the generation of reactive oxygen species (ROS) by NADPH oxidase and degranulation of antimicrobial proteins/peptides, such as myeloperoxidase (MPO) and neutrophil elastase (NE), into the phagosome and extracellular space^14,15^. In addition, neutrophils can release extracellular web-like structures, known as neutrophil extracellular traps (NETs), made of decondensed DNA and coated with antimicrobial effectors^16–18^. NETs are released during NETosis, a programmed cell death requiring ROS production and the granule proteins MPO and NE^19,20^. Upon activation, MPO facilitates the translocation of NE from azurophilic granules to the nucleus, where it cleaves histones to decondense chromatin^21,22^. Together, these effector functions kill extracellular Gram-negative and Gram-positive bacterial pathogens^16–18,23^.

Bacterial cell wall components are critical activators of innate immune responses. In Gram-negative bacteria, lipopolysaccharides (LPS) are a key component of the outer membrane, essential for maintaining structural integrity and facilitating host interactions^24,25^. LPS is composed of three regions, each with a distinct role: lipid A, which anchors LPS to the bacterial membrane and acts as an endotoxin; the core oligosaccharide, which links lipid A to the O-antigen (OA) and maintains membrane stability, while also contributing to immune evasion; and the OA, a variable polysaccharide chain that extends outward, helping the bacteria evade immune recognition by masking conserved bacterial structures^24–27^. LPS is a potent activator of the innate immune system, triggering inflammation through various pathways. Specifically, lipid A is recognized by immune receptors such as TLR4 and caspase 4/11^26–32^. This recognition by neutrophils leads to the production of reactive oxygen species (ROS), degranulation, and NETs formation, all crucial for bacterial clearance^33–35^. Meanwhile, the variability of the OA aids in immune evasion by allowing the bacteria to alter its surface structures, thereby avoiding detection by the host’s immune system^36,37^. Together, these components enable LPS to serve both as a structural feature and as a key factor in host-pathogen interactions.

Despite the important role neutrophils play in the detection and elimination of Gram-negative pathogens, little is known about their role in combating hvKp infection. A few studies have described the role of LPS and the hypervirulent capsule in protecting against neutrophil phagocytosis and bactericidal activity, as opposed to cKp isolates^10,38–44^. However, the underpinning mechanism remains unknown. Recent findings suggested that excessive ethanol produced by high-alcohol-producing Kp strains inhibits neutrophil phagocytosis and immune responses, increasing susceptibility to Kp^45^.

In this study, using a pulmonary mouse infection model, we show that while neutrophils are recruited in large numbers to the lung, they cannot control hvKp infection. Using ex vivo assays, we demonstrated that the hypervirulent capsule functions as a protective barrier, preventing bacterial recognition and neutrophil activation, thereby blocking ROS production, degranulation, and NET formation. In isogenic acapsular (Δ*wcaj*) or hypermucoid capsule determinant (Δ*rmpADC*) hvKp mutants, recognition of the OA was essential to specifically trigger neutrophil degranulation and bacterial killing, although OA recognition was not required for NET formation. In the absence of OA, neutrophils were more efficient in killing acapsular hvKp, suggesting that OA helps the bacteria to evade NET-mediated destruction. Taken together, our findings highlight the important role of the hypervirulent capsule as a shield, preventing neutrophil activation and reveal a dual function for OA, which function as a strong neutrophil activator while also protecting the hvKp from NET-mediated killing.

## Results

### hvKp survives in infected lungs despite robust neutrophil recruitment

The aim of this study was to investigate the mechanism underpinning hvKp virulence. To this end, we analysed lung immune responses following the intratracheal (IT) inoculation of 500 CFU of the prototype hvKp strain ICC8001 (K2/O1; a derivative of ATCC43816) (**S. Table 1**) into CD1 mice. At 24, 48 and 72 h post infection (hpi), we collected lung homogenate, bronchoalveolar lavage (BAL) and blood from hvKp ICC8001-infected mice; mock-infected (PBS) mice were used as controls. At 72 hpi, infected mice developed significant weight loss compared to controls (**Fig. 1A**). Quantification of the bacterial burden revealed temporal increases in hvKp ICC8001 CFU counts in both the lung and blood (**Fig. 1B**).

**Figure 1.**
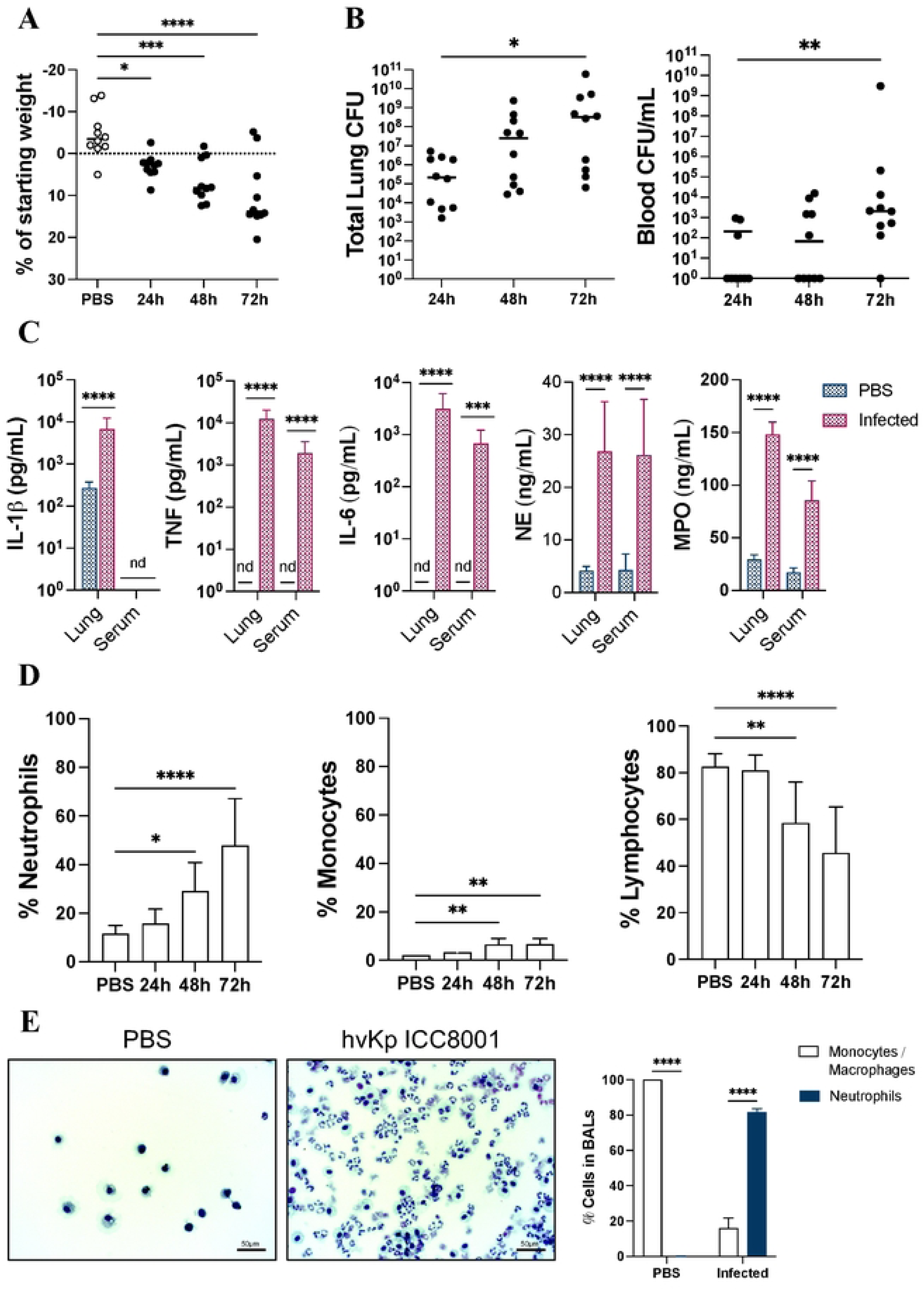
hvKp ICC8001 replicates in infected lungs. (**A**) Percentage of weight loss at 24, 48 and 72 hpi, relative to starting weight (day 0). (**B**) hvKp ICC8001 load in the lungs and blood at 24, 48 and 72 hpi. (**C**) ELISA of IL-1β, IL-6, TNF, NE and MPO from serum and lung homogenates of uninfected and infected mice at 72 hpi. (**D**) Percentage of neutrophils, monocytes and lymphocytes in the blood of infected mice at 24, 48 and 72 hpi. (**E**) Representative images of BAL from PBS and infected mice at 72 hpi, using Kwik-Diff staining. Scale bars, 50µM. The percentage of neutrophils (poly-lobulated nucleus) and monocytes/macrophages (mono-lobulated nucleus) calculated relative to total amount of cells per image. All experiments were conducted in 2 biological replicates with 5 mice per group. (**A, C-E**) Significance was determined by ordinary one-way ANOVA or (**C**) two-way ANOVA, followed by Dunnett’s and Sidak’s multiple-comparison post-test, respectively. (**B**) Significance was determined by Mann-Whitney T-test. (**A-D**) All comparisons not shown were non-significant. (**A-B**) Values are expressed as mean or (**C-D**) as mean +/- SEM. *p< 0.05, **p< 0.01, ***p< 0.001 ****p< 0.0001.

While hvKp ICC8001 infection led to increased levels of IL-1b specifically in the lung homogenate (**Fig. 1C**), the levels of interleukin-6 (IL-6) and TNF, as well as myeloperoxidase (MPO) and neutrophil elastase (NE), increased in both the lung homogenate and serum (**Fig. 1C**). This was associated with a significant increase in blood neutrophils, along with a smaller increase in blood monocytes, while lymphocyte counts significantly decreased (**Fig. 1D**). Recruited neutrophils account for approximately 80% of the white blood cells in BAL, few of which contained intracellular hvKp ICC8001 (**Figs. 1E, S1A**). No neutrophils were seen in the BAL of mock infected mice (**Fig. 1E**). These results suggest that despite extensive recruitment, neutrophils were unable to kill hvKp ICC8001. The processes implicated in neutrophil activation and bacterial killing are shown schematically in **Fig. 2A**.

**Figure 2.**
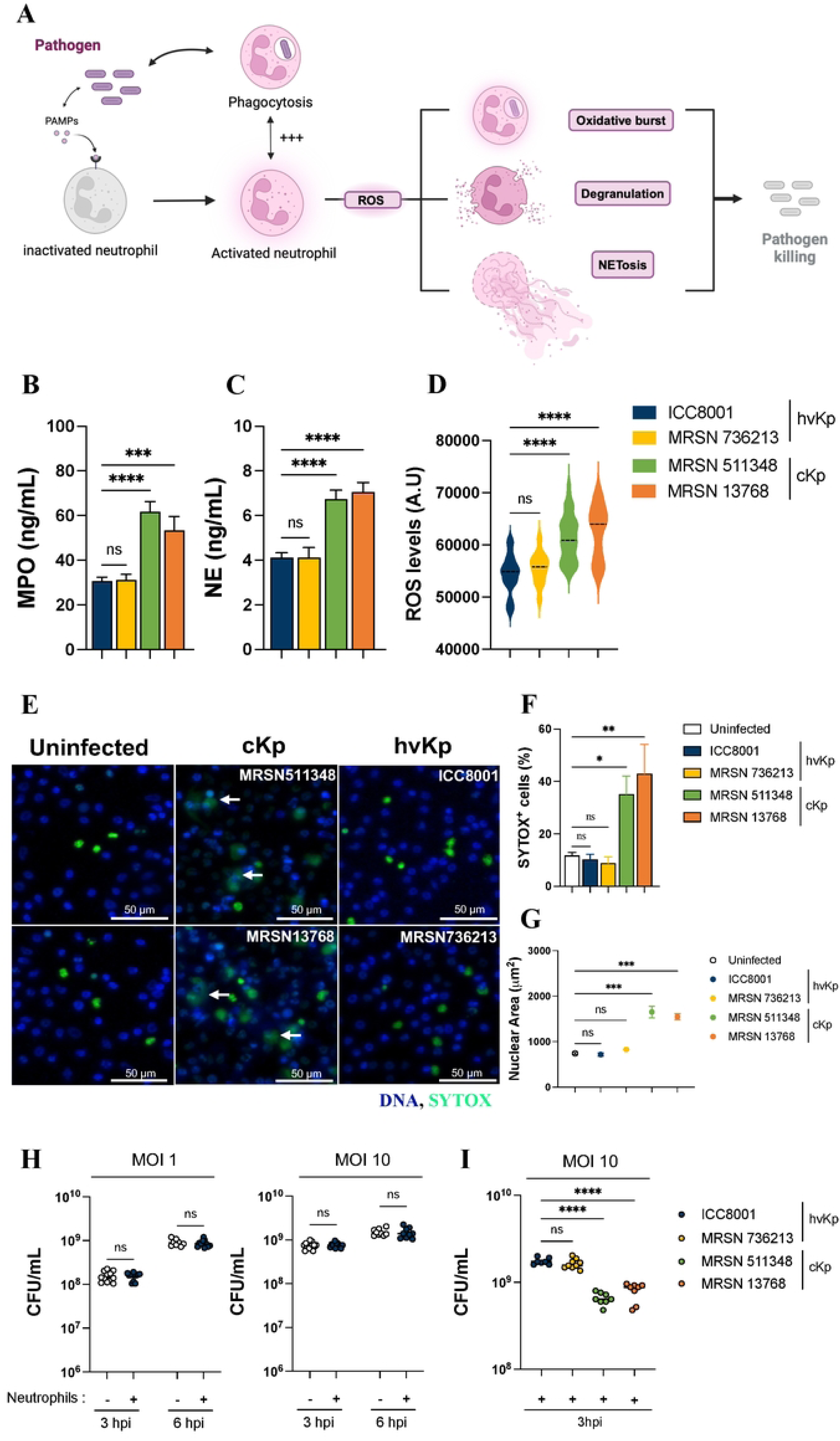
hvKp ICC8001 dampens neutrophil activation and killing. (**A**) Schematic of neutrophil activation. (**B-D**) Levels of MPO (**B**), NE (**C**) and ROS (**D**) in the supernatant of BMDNs infected for 3 h with different cKp and hvKp strains. (**E**) DNA straining of BMDNs infected with different hvKp and cKp strains, using SYTOX and DAPI. Scale bars, 50µM. Representative images of three independent experiments. (**F**) Quantification of SYTOX^+^ neutrophils. (**G**) Quantification of the neutrophils nuclear area as a measure of NETs. (**H**) Total hvKp ICC8001 load in BMDNs, at 3-6 hpi (MOI 1 and 10). (**I**) Total hvKp and cKp load in BMDNs, 3 hpi (MOI 10). (**B-D, F-I**) Significance to hvKp ICC8001 was determined by ordinary one-way ANOVA, followed by Dunnett’s multiple-comparison post-test. Values are expressed as (**B-D, F-G**) mean +/- SEM or **(H-I)** mean. ns (no significant) p>0.05, *p< 0.05, **p< 0.01, ***p< 0.001, ****p< 0.0001. Data are representative of three independent experiments.

### hvKp dampens neutrophil activation and killing

To identify the bacterial factors protecting hvKp ICC8001 from clearance by neutrophils in vivo, we set up functional ex vivo assays using bone marrow derived neutrophils (BMDNs), isolated from C57BL/6 mice. We first investigated if LPS priming affected BMDNs responses to hvKp infection. Comparing naïve and primed BMDNs, with and without PMA-stimulation, revealed greater MPO/NE release and ROS production following priming (**Figs. S2B, S2C)**.

However, infection of both naïve and LPS-primed BMDNs with hvKp ICC8001 did not trigger more release of NE, MPO or ROS compared to uninfected BMDNs (**Fig. S2B)**. We therefore mainly used un-primed BMDNs for the ex vivo functional assays; unless mentioned otherwise, MOI of 10 was used for infection.

We infected BMDNs with either HMV^+^ hvKp strains ICC8001 or MRSN_736213 (K2/01, clinical isolate) or with HMV^-^ cKp strains MRSN_511348 (K2/O1) and MRSN_13768 (K17/O1), both clinical isolates (**Figs. S2A, S. Table 1**). Infection with the cKp strains MRSN_511348 and MRSN_13768 tiggered significantly greater release of MPO (**Fig. 2B**), NE (**Fig. 2C**) and ROS (**Fig. 2D**) compared to infection with the hvKp strains ICC8001 and MRSN_736213.

We next assessed the ability of BMDN to produce NETs in response to hvKp or cKp infections, using the cell-impermeable SYTOX dye, which only stains extracellular DNA or DNA in permeabilised (dying) cells. While sporadic cell death with condensed nuclei was seen in the uninfected negative control BMDNs (**Figs. 2E, S3A, S3B**), infection of BMDNs with the cKp strains induced higher SYTOX incorporation over time (**Fig. S3B)**, associated with decondensed nuclei and extracellular DNA (**Figs. 2E** white arrows, **S3A).** In contrast, BMDNs infected with the hvKp strains led to SYTOX staining similar to the uninfected control cells, associated with low nuclear areas (**Figs. 2E, S3A, S3B**). Quantification revealed significantly greater number of SYTOX^+^ neutrophils, with increased DNA area, following infection with the cKp compared to hvKp strains (**Fig. 2F, 2G**). Moreover, infection of BMDNs with either multiplicity of infection (MOI) of 1 or 10 revealed that hvKp ICC8001 grew equally in the presence of un-primed or LPS-primed neutrophils (**Figs. 2H, S3C)**. In contrast, neutrophils were effective in killing the cKp strains MRSN_511348 and MRSN_13768 (**Fig. 2I**).

We then investigated the ability of BMDN to phagocytose hvKp. This revealed that >99% of the hvKp ICC8001 remained extracellular, with only 0.2% found to be intracellular at 6 hpi (**Figs. 3A, 3B**). Consistent with the lack of MPO, NE, ROS and NET release, hvKp ICC8001 proliferated both intra- and extra-cellularly over time in the presence of BMDNs (**Figs. 3A, 3B)**. Transmission electron microscopy revealed a higher number of hvKp ICC8001 at 6 compared to 3 hpi, within specious vacuoles (**Fig. 3C**). Notably, the increase in intracellular bacteria was primarily due to proliferation rather than enhanced phagocytosis, as the addition of gentamicin at 2.5 or 5.5 hpi resulted in similar number of intracellular bacteria at 6 hpi (**Fig. 3D**). Together, these results suggest that expression of hypervirulence genes prevent neutrophil responses to hvKp infection, which can survive and grow in their presence.

**Figure 3.**
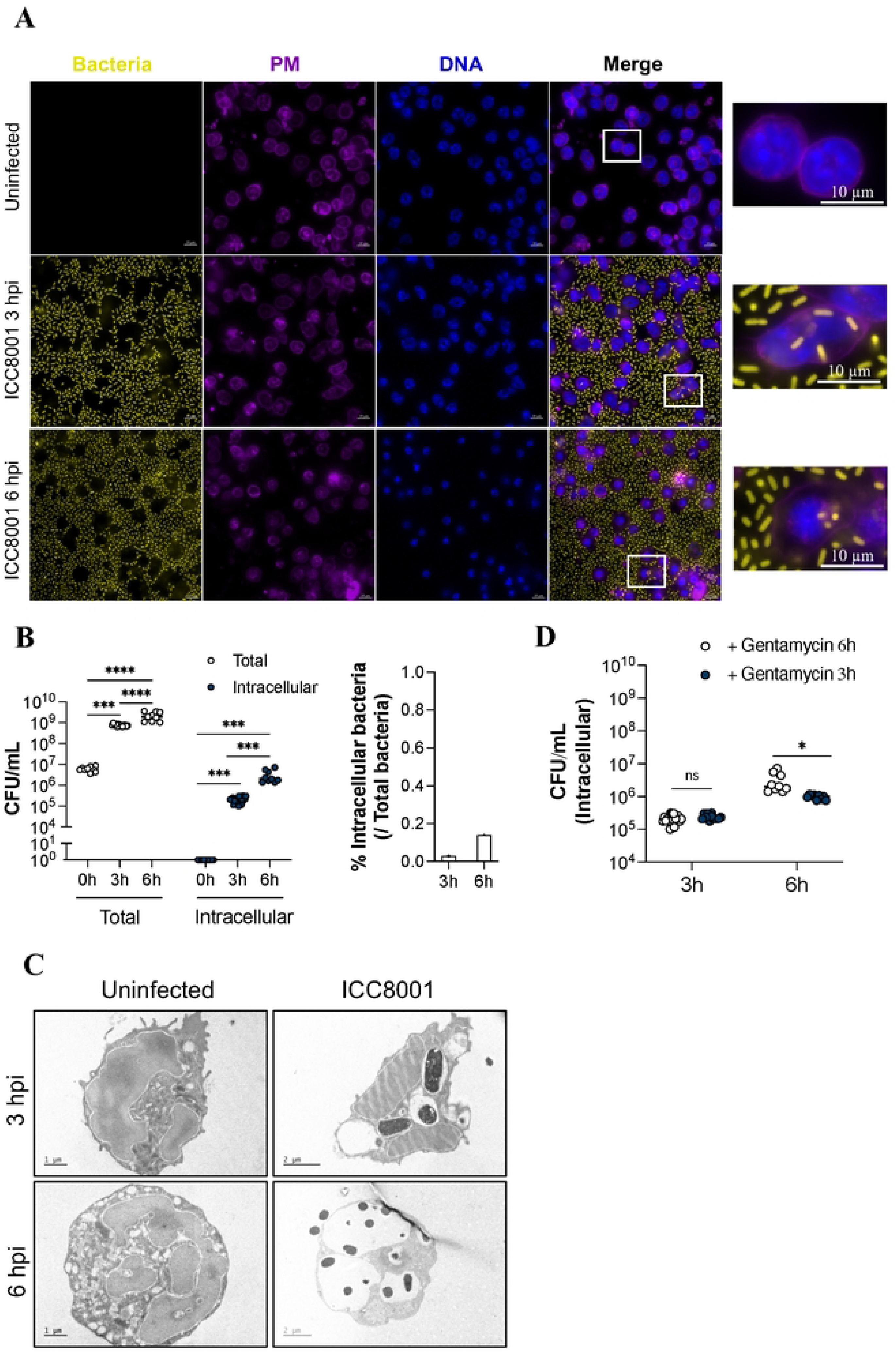
hvKp ICC8001 escapes neutrophil phagocytosis and killing. (**A**) hvKp ICC8001 (yellow) localisation in BMDNs at 3-6 hpi. WGA (purple) and DAPI (blue) were used to stain neutrophils. Scale bars, 10µM. (**B**) Total and intracellular hvKp load in BMDNs at 3h and 6 hpi. (**C**) Representative transmission electron microscopy (TEM) images of intracellular hvKp in uninfected or infected BMDNs at 3 and 6 hpi. Scale bars, 1&2µM. (**D**) Intracellular hvKp load in BMDNs, at 3h and 6 hpi, in the presence of gentamycin, added at 2.5 or 5.5 hpi. (**B, D**) Significance was determined by ordinary two-way ANOVA, followed by Tukey’s (**B)** and Sidak’s **(D**) multiple-comparison post-test. Values are expressed as mean. ns (not significant) p>0.5, *p< 0.05, ***p< 0.001, ****p< 0.0001. Data are representative of three independent experiments.

### The hypervirulent capsule protects hvKp ICC8001 from neutrophil phagocytosis and killing

Given that hvKp strains are characterised by hypercapsulation and HMV, we next investigated the role of the hypervirulent capsule in evasion of neutrophil responses. To this end, BMDNs were infected with hvKp ICC8001, a capsule-deficient mutant Δ*wcaJ* or a mutant missing the regulators resulting in hypercapsulation and HMV (Δ*rmpADC)* **(Figs. S2A, S. Table 1)**. The absence of either the capsule or *rmpADC* resulted in increased phagocytosis and enhanced intra- and extracellular killing **(Figs. 4A-C)**.

**Figure 4.**
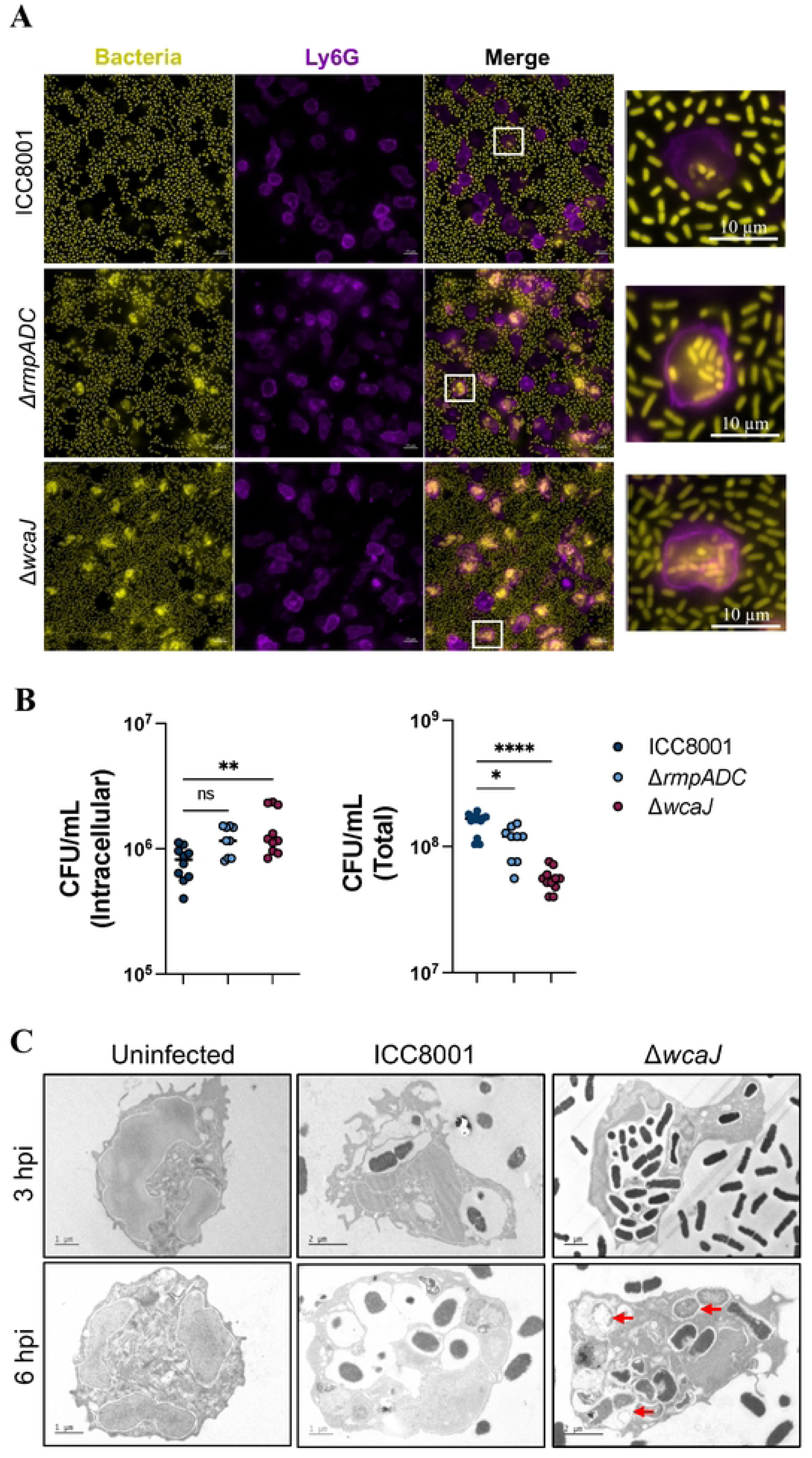
The hypervirulent capsule protects hvKp ICC8001 from neutrophil phagocytosis and killing. (**A**) Localisation of hvKp ICC8001 (yellow), Δ*wcaJ* and Δ*rmpADC* in BMDNs at 3 hpi. Ly6G (purple) and DAPI (blue) were used to stain neutrophils. Scale bars, 10µM. (**B**) Total and intracellular hvKp ICC8001, Δ*wcaJ* and Δ*rmpADC* load in BMDNs, at 3 hpi. (**C**) TEM images of intracellular hvKp ICC8001 or Δ*wcaJ* in BMDNs at 3 and 6 hpi. Scale bars, 1-2µM. Red arrows indicate intracellular bacteria killing. (**B**) Significance to hvKp ICC8001 was determined by ordinary one-way ANOVA, followed by Dunnett’s multiple-comparison post-test. Values are expressed as mean. ns (not significant) p>0.05, *p< 0.05, **p< 0.01, ****p< 0.0001. Data are representative of three independent experiments.

We hypothesised that the presence of a hypervirulent capsule could protect hvKp from neutrophil recognition and subsequent activation. To test this hypothesis, we quantified ROS production and NE and MPO localisation in BMDNs infected with either hvKp ICC8001, Δ*wcaJ* or Δ*rmpADC.* The absence of capsule or *rmpADC* induced a burst of ROS (**Fig. 5A**) and increased release of NE and MPO into the extracellular space (**Fig. 5B**), which was comparable to cKp strains MRSN_511348 and MRSN_13768 infection (**Fig. 5C**); PMA stimulation and actin were used as a positive and loading controls (**Fig. 5B**). Moreover, NE and MPO translocation to the nucleus was only observed in the absence of capsule, while they remained in the granules following hvKp ICC8001 infection (**Figs. 5D, S4A)**. Ultimately, the absence of capsule was associated to NET formation and rendered hvKp ICC8001 susceptible to neutrophil killing, comparable to cKp (**Figs. 5E, 5F, S4B**).

**Figure 5.**
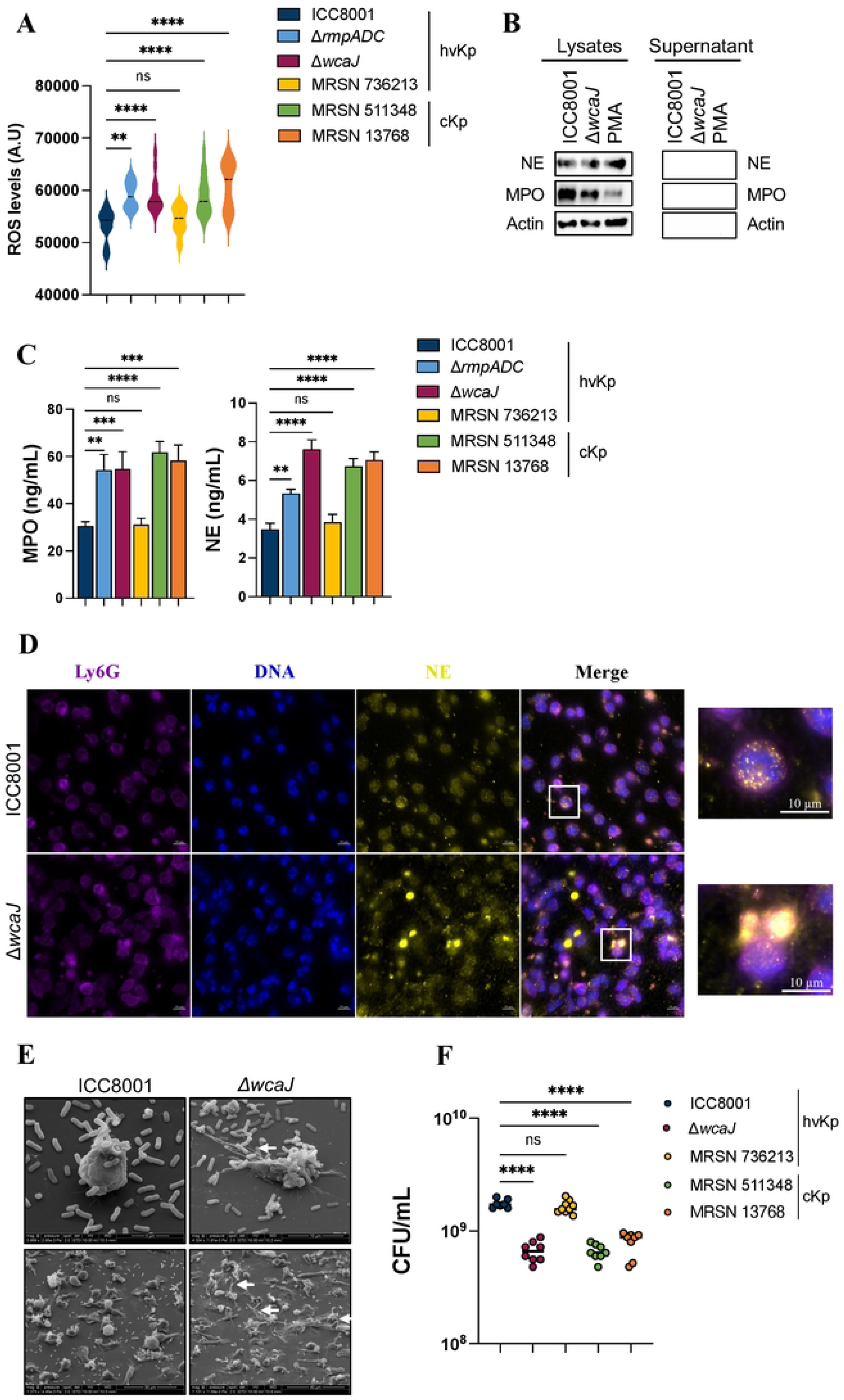
The hypervirulent capsule prevents neutrophil activation. (**A**) ROS levels in BMDNs, infected with different hvKp or cKp strains for 3 h. (**B**) Immunoblot analysis of MPO and NE release from BMDNs stimulated with PMA or infected with hvKp ICC8001 or Δ*wcaJ* at 3 hpi. Actin was used as loading control. Data are representative of three independent experiments. (**C**) Levels of NE and MPO in the supernatant of BMDNs, infected for 3 h with different cKp and hvKp strains. (**D**) Localisation of NE (yellow) in BMDNs infected with hvKp ICC8001 or Δ*wcaJ* for 3 h. Ly6G (purple) and DAPI (blue) were used to stain neutrophils. Scale bars, 10µM. (**E**) Scanning Electron Microscopy (SEM) images of BMDNs infected with hvKp ICC8001 and *ΔwcaJ* at 3 hpi. Arrows point at NETs. Scale bars, 5, 10, 30 and 40µM. (**F**) Total hvKp and cKp strains load in BMDNs at 3 hpi. (**A, C, F**) Significance to hvKp ICC8001 was determined by ordinary one-way ANOVA, followed by Dunnett’s multiple-comparison post-test. (**A, C**) Values are expressed as mean +/- SEM or (**F**) mean. ns (not significant) p>0.05, **p< 0.01, ***p< 0.001, ****p< 0.0001. Data are representative of three independent experiments.

To investigate whether NETosis alone was responsible for controlling acapsular hvKp, we infected BMDNs isolated from MPO-deficient mice (which are unable to form NETs)^22^ with hvKp ICC8001 or Δ*wcaJ*. While similar hvKp ICC8001 CFUs/ml were recovered from infected WT and MPO-deficient BMDNs, we observed significantly improved killing of Δ*wcaJ* in the absence of MPO, suggesting that NETs are not required for clearance (**Fig. 6A**). The improved killing in MPO-deficient neutrophils was associated with increase NE release into the extracellular space as well as intracellular killing (**Figs. 6B, 6C**). We hypothesised that, while MPO is required for NE translocation from the granules into the nucleus leading to NETs formation, MPO is not needed for NE delivery into the phagosome or extracellular space. Hence, NE activity in these compartments, along with potentially other granule-derived proteases, could be crucial for Δ*wcaJ* and cKp killing. To test this hypothesis, we inhibited the serine proteases stored in the granules, including cathepsin G, NE and proteinase 3, using the irreversible serine protease inhibitor, diisopropyl fluorophosphate (DFP). While the serine protease inhibitor did not affect killing of the hvKp ICC8001 control, it protected the Δ*wcaJ* and cKp strains from neutrophil killing (**Fig. 6D)**. Overall, these data suggest that in the absence of a hypervirulent capsule, serine proteases released from granules following neutrophil activation are key to effectively clear cKp and acapsular hvKp.

**Figure 6.**
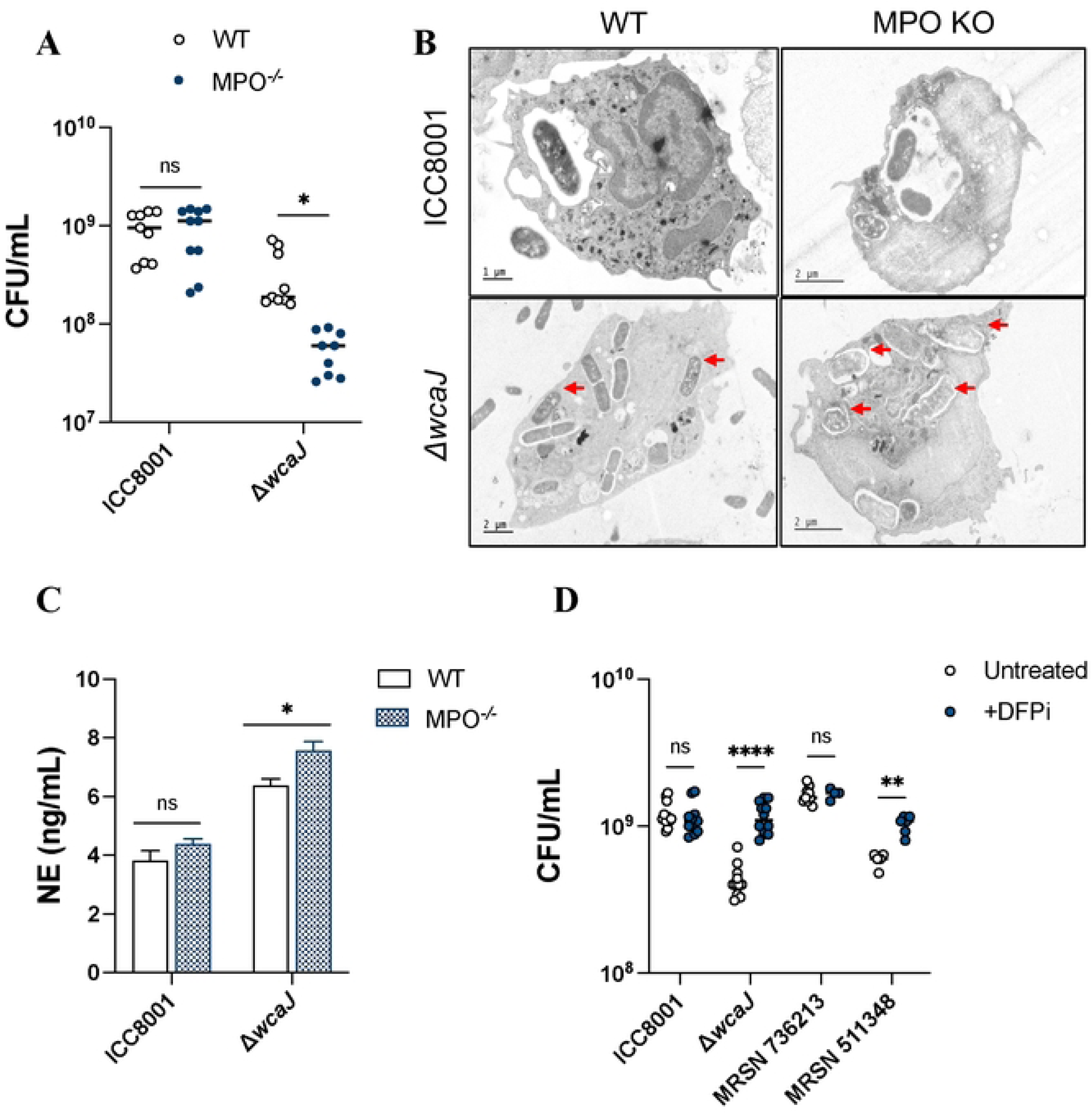
Serine proteases effectively kill acapsular hvKp and cKp. (**A**) Total hvKp ICC8001 and Δ*wcaJ* load in BMDNs from WT or MPO^-/-^ mice, 3 hpi. (**B**) TEM images of intracellular hvKp WT or Δ*wcaJ,* in BMDNs from WT or MPO^-/-^ mice, 3 hpi. Scale bars, 1 and 2µM. Red arrows indicate intracellular bacteria killing. (**C**) Levels of NE in the supernatant of BMDNs from WT or MPO^-/-^ mice, infected for 3 h with hvKp strains ICC8001 and Δ*wcaJ.* (**D**) Total hvKp and cKp strains load in untreated or DFP-treated BMDNs, 3 hpi. (**A, C)** Significance was determined by multiple Mann-Whitney tests. (**D)** Significance was determined by ordinary two-way ANOVA, followed by Sidak’s multiple-comparison post-test. (**A, D**) Values are expressed as mean or (**C**) mean +/- SEM. ns (not significant) p>0.05, *p< 0.05, **p< 0.01, ****p< 0.0001. Data are representative of three independent experiments.

### The OA is needed for neutrophil degranulation but dispensable for NETs formation

We next aimed to identify the virulence factor leading to neutrophil activation in the absence of the hypervirulent capsule. We hypothesised that in its absence, LPS-particularly the exposed OA region could drive neutrophil activation and subsequent bacterial killing. To test this, bacterial killing and neutrophil responses were assessed in BMDNs infected with either hvKp ICC8001, Δ*wcaJ,* Δ*rfb* (which expresses capsule, lipid A and core region of LPS but lacks OA, known as ‘rough’) or the Δ*rfb /*Δ*wcaJ* double mutant, which is acapsular and rough. Infection revealed that while the absence of OA in Δ*rfb* did not trigger neutrophil degranulation, the enhanced release of MPO and NE triggered by Δ*wcaJ* was abolished in the absence of the OA (Δ*rfb /*Δ*wcaJ*) (**Fig. 7A**). Interestingly, while OA is needed for neutrophil degranulation, it is not required for NETs formation as, despite a decrease in the number of SYTOX-positive cells, a similar percentage of neutrophils still undergo NETosis (**Figs. 7B-D**). Moreover, while the absence of OA alone did not affect hvKp ICC8001 survival, Δ*wcaJ,* and to a greater extent Δ*rfb/*Δ*wcaJ,* were more efficiently killed by the BMDNs (**Fig. 7E)**. These findings indicate that recognition of the OA specifically triggers neutrophil degranulation during infection with acapsular hvKp ICC8001, although it is not essential for NET formation (**Fig. 7F)**. This underscores the idea that different stimuli and pathogen types can provoke distinct neutrophil responses, enabling a targeted defence that prevents unnecessary widespread inflammation. Additionally, the results suggest that OA may act as an extra protective layer, helping the bacteria evade NETs and enhance their survival.

**Figure 7.**
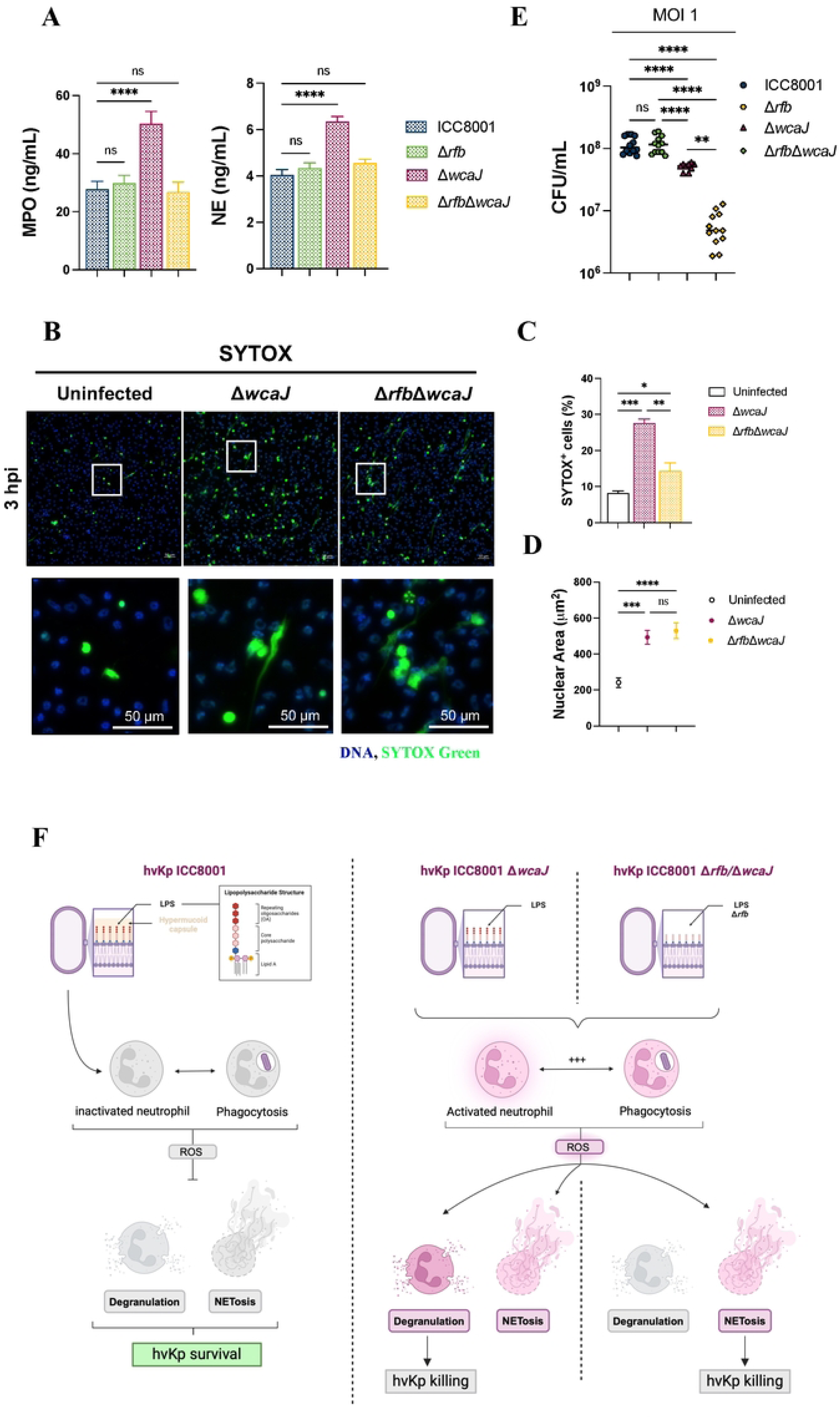
The OA is needed for neutrophil degranulation but dispensable for NET formation. (**A**) Levels of NE and MPO in the supernatant of BMDNs at 3 hpi with hvKp ICC8001, Δ*wcaJ,* Δ*rfb* or Δ*rfb*Δ*wcaJ*. (**B**) DNA straining of BMDNs infected for 3 h with hvKp ICC8001, Δ*wcaJ*, or Δ*rfb*Δ*wcaJ*, using SYTOX and DAPI. Scale bars, 50µM. Representative images of three independent experiments. (**C**) Quantification of SYTOX+ neutrophils. (**D**) Quantification of the neutrophils nuclear area as a measure of NETs. (**E**) Total hvKp ICC8001, Δ*wcaJ,* Δ*rfb*, or Δ*rfb*Δ*wcaJ* load in BMDNs at 3 hpi. (**F**) Schematic of hvKp infection in BMDNs. (**A**) Significance to hvKp ICC8001 was determined by ordinary one-way ANOVA, followed by Dunnett’s multiple-comparison post-test. (**C-E)** Significance was determined by ordinary one-way ANOVA, followed by (**C-D)** Dunnett’s multiple-comparison post-test or (**E)** Tukey’s multiple-comparison post-test. (**A, C-D**) Values are expressed as mean +/- SEM (**E**) or mean. ns (not significant) p>0.5, *p< 0.05, **p< 0.01, ***p< 0.001, ****p< 0.0001. Data are representative of three independent experiments.

## Discussion

Neutrophils play a vital role in the immune response against pathogens. Their activation relies on the recognition of pathogen-associated molecular patterns (PAMPs) such as lipid A of LPS, activating several inflammatory and killing processes including phagocytosis, cytokine release, ROS production, degranulation, and the formation of NETs^14,46^. Impaired neutrophil function can lead to increased susceptibility to infections, highlighting their critical role in host defense^14,16–18^.

While the capsule has long been known to enhance virulence of hvKp, the molecular and cellular basis underpinning this was not known. In this study, we have shown that the hypervirulent capsule of hvKp strains enables the bacteria to evade neutrophil recognition and killing. In vivo, hvKp infection led to significant neutrophil recruitment to the lungs but resulted in no effective bacterial clearance. Mechanistically, ex vivo experiments demonstrated that the hypervirulent capsule not only acts as a protective barrier against phagocytosis but also conceals the OA, preventing neutrophil degranulation. While the lipid A and inner core LPS components are a well-studied PAMPs, implicated in activation of sensors like TLR4^28,33,34^, caspase-4 and GBP2^30–32^, we now show that the Kp OA itself is a PAMP that activates degranulation; the sensor of which in neutrophils remains unknown.

Our findings also indicate that while the presence of OA is essential for inducing neutrophil degranulation - evidenced by the release of MPO and NE - its recognition is not required for NET formation. This highlights a unique aspect of the neutrophil response where Kp OA acts as a signal, triggering specific defensive actions without orchestrating a broader inflammatory response. Previous work has already described a model in which neutrophil phagocytosis acts as a checkpoint to dictate the decision to form NETs^46^. This distinction suggests that the activation pathways may be compartmentalised or limited in neutrophils. While different stimuli and sensors can induce degranulation and/or NETosis, the specific immune landscape (e.g. cytokines, DAMP), type of pathogen, and signalling pathways involved can determine which response predominates. Both processes play crucial roles in the immune response, and their regulation is vital for maintaining a balanced response to infections and preventing tissue damage. Thus, by concealing OA within a hypervirulent capsule, hvKp can successfully evade neutrophil detection and subsequent activation, contributing to its pathogenicity.

Interestingly, the absence of OA also correlated with improved bacterial killing by NETs, indicating that OA might also confer protective benefits to the bacteria, allowing them to withstand neutrophil attacks more effectively. These findings highlight the dual function of OA: it not only serves as a recognition signal that activates neutrophils but also appears to play a role in protecting the bacteria from NETs.

We have shown that like the *rmpADC* deletion mutant of hvKp ICC8001, clinical cKp strains (both expressing O1) also trigger neutrophil degranulation. This suggests that the capsule in cKp does not mask the OA, providing a mechanistic explanation for the Kp epidemiology, i.e. by the virtue of their thick capsule, which masks OA, hvKp are more pathogenic as they evade neutrophil activation and are thus capable of infecting immunocompetent individuals.

Overall, our findings contribute new insights into the mechanisms of immune evasion by hvKp and emphasize the importance of neutrophil “decision-making” in pathogen clearance. Targeting these pathways might be a promising avenue for improving host defence against this and potentially other encapsulated pathogens.

## Acknowledgments

We would like to thank Venizelos Papayannopoulos and Drinalda Cela from the Crick Institute (London), for the MPO^-/-^ neutrophils; Wen Wen Low for the generation of the Δ*wcaJ* mutant; Renaud Poincloux from IPBS (Toulouse), for the Scanning Electron Microscopy images and the METi imaging facility (CBI, Toulouse), member of the national infrastructure France-BioImaging supported by French National Research Agency (ANR-10-INBS-04) for the Transmission Electron Microscopy images. We also want to thank thank Venizelos Papayannopoulos for critical reading of the manuscript. This project was supported by a Wellcome Trust Investigator Award to G.F (224282/Z/21/Z)).

## Materials and Methods

### Ethics statement

All animal work was conducted under the auspices of the United Kingdom Animals (Scientific Procedures) Act 1986 (License: PP7392693) and was locally approved by the institutional ethics committee (the Animal Welfare and Ethical Review Body [AWERB]).

### Mice infections

Female CD1 mice, 28-30g weight (Charles River, UK), were randomised into groups (co-housed in groups of 5) with a one-week acclimatization period prior to infection. The mice were kept on a 12-hour light-dark cycle and had ad libitum access to food and water.

Inoculum was prepared from overnight cultures grown in Luria Broth (LB), diluted in 1xPBS to a total volume of 50µL, and delivered via intratracheal intubation under ketamine (80mg/kg) and medetomidine (o.8mg/kg) anaesthesia. Recovery from anaesthesia was monitored at 32-35°C following the administration of atipamezole (1mg/kg) reversal. Inoculum was verified (+/- 10%) by enumeration on agar plates.

Animals were sacrificed at indicated times after infection and bronchoalveolar fluids (BALFs), blood and lungs were recovered. Bacterial enumeration (CFU) was conducted on LB agar plates supplemented with 50µg/ml rifampicin. Blood samples were collected by cardiac puncture under terminal anaesthesia (ketamine 100mg/kg, medetomidine 1mg/kg). BALFs were recovered under terminal anaesthesia (pentobarbital 200 ml of 200 mg/ml, i.p.) in 3mL of 1X PBS, supplemented with 3mM of EDTA (Invitrogen, 11568896). Lungs were excised post-mortem and homogenized. Samples were subjected to 10-fold serial-dilution before plating on agar plates and incubated overnight at 37°C.

### Generation of hvKp mutants

All genomic mutations were made in ICC8001, a rifampicin-resistant derivative of *K. pneumoniae* ATCC43816 using a two-step recombination methodology, as described^47^. Strains used are listed in supplementary Table 1.

### Bacterial Cultures

*K. pneumoniae* strains were grown overnight in LB medium, containing 50µg/mL of streptomycin, at 37°C with constant agitation (200 rpm), and sub-cultured the next day by diluting overnight culture 1:100 at 37°C, with constant agitation (200 rpm), until reaching an optical density (OD) of 1. All *K. pneumoniae* strains used in the study are listed in supplementary Table 1.

### Isolation of primary murine neutrophils

Murine bone marrow cells were isolated from tibias and femurs of C57BL/6 mice. Neutrophils were purified by positive selection using Anti-Ly-6G MicroBead Kit (Miltenyi Biotech) according to manufacturer’s instructions. This process routinely yielded >95% of neutrophil population as assessed by flow cytometry of Ly6G^+^/CD11b^+^ cells.

### Cell plating and infection of neutrophils

Following isolation, neutrophils were centrifugated for 10 min at 300xg and pellet was resuspended in serum free Opti-MEM medium (Gibco). Absolute cell number was determined with automated countess 3 cell counter (Invitrogen) with trypan blue cell death exclusion method (typically living cells represent > 80% of cell solution) and cell density was adjusted at 10^6^ / mL by adding Opti-MEM culture medium. Neutrophils were then plated in either 96 well plates, 24 well plates or 6 well plates with 50 µL (5.10^4^ cells), 500 µL (5.10^5^ cells) or 2 mL (2.10^6^ cells) respectively. When indicated, cells were primed with LPS (200 ng/ml) for 3 hours. Neutrophils were infected with various bacterial strains and multiplicity of infections (MOI) as indicated in figure legends, centrifuged for 5 min at 700xg and incubated at 37°C in a 5% carbon dioxide (CO_2_) incubator the indicated time.

### Kinetic analysis of SYTOX Green incorporation assay

Cells were plated at density of 5.10^4^ cells per well in Black/Clear 96-well Plates (Nunc) in Opti-MEM culture medium supplemented with SYTOX-Green dye (100ng/mL) (ThermoFisher, S7020) and infected/treated as mentioned in figure legends. Green fluorescence was measured in real-time using FLUOstar Omega plate reader (BMG Labtech) at 37°C with 5% CO_2_. Maximal cell death was determined with whole cell lysates from unstimulated cells incubated with 0.1% Triton X-100.

### ELISA(s)

Mouse IL-6 (ThermoFisher, 88-7064-88), IL-1β (ThermoFisher, 88-7324-88), TNF (ThermoFisher, 88-7013A-88), NE (R&D, DY4517) and MPO (R&D, DY3667) levels were quantified according to the manufacturer’s instructions.

### Quantification of ROS levels

Cells were plated at density of 5.10^4^ cells per well in Black/Clear 96-well Plates (Nunc) in Opti-MEM culture medium supplemented with the cell-permeant 2’,7’-dichlorodihydrofluorescein diacetate (H2DCFDA) (10µM) (ThermoFisher, D399) and infected/treated as mentioned in figure legends. Fluorescence was measured in real-time using FLUOstar Omega plate reader (BMG Labtech) at 37°C with 5% CO_2_.

### Quantification of CFU loads

Neutrophils were plated at a density of 5.10^5^ cells per well in a 24-well plate in Opti-MEM culture medium and infected with different Kp strains at an MOI of 1 or 10 for 3 to 6h. At the end of the infection, total bacterial load was determined by lysing cells with 0.1% Triton X-100 for 10 min and subjected to 10-fold serial-dilution before plating on agar plates and incubated overnight at 37°C. Phagocytosis assay was performed by adding gentamicin (100µg/mL) for 45 min, 2h post-infection, washing 3 times the cells before adding fresh Opti-MEM medium with 0.1% Triton X-100. Intracellular CFU counts was determined by performing a 10-fold serial-dilution before plating on agar plates and incubated overnight at 37°C.

### Immunoblot analysis

Neutrophils were plated at a density of 2.10^6^ cells per well in 6 well plates and infected with different Kp strains at an MOI of 10 for 3h. At the end of the infection, 5 mM of diisopropylfluorophosphate (DFP) cell permeable serine protease inhibitor (Sigma, D0879) was added to cell culture medium. Cell supernatant was collected and clarified from non-adherent cells by centrifugation for 10 min at 300xg. Cell pellet and adherent cells were lysed in 100μL of RIPA buffer (Sigma, R0278) supplemented with 5 mM diisopropylfluorophosphate (DFP) in addition to the protease inhibitor cocktail (Sigma, 4693159001). Cell scrapper was used to ensure optimal recovery of cell lysate. Collected cell lysate was homogenized and supplemented with 1X laemli buffer (Bio-rad, 1610747) before boiling samples for 10 min at 95 °C. Soluble proteins from cell supernatant fraction were precipitated as described previously^48^. Precipitated pellet was then resuspended in 100μL of RIPA buffer plus laemli, supplemented with 5 mM of DFP and protease inhibitor cocktail, and heated for 10 min at 95 °C. Samples were resolved by SDS-PAGE and transferred to PVDF membranes. Cell lysate and supernatant fraction were analysed individually. Non-specific binding was blocked with 5% skim milk (Sigma, 70166) and membrane were probe (1:100) with anti-MPO (abcam, ab208670), anti-NE (abcam, ab314916) and anti-ß-Actin (Cell signalling, 3700S). Primary antibodies were detected with horseradish peroxidase-conjugated secondary antibodies goat anti-rabbit (Advansta, R-05072-500) and goat-anti-mouse (Advansta, R-05071-500).

### Immunofluorescence

Neutrophils were plated at a density of 5.10^5^ cells on glass bottom Ibidi chambered coverslips in Opti-MEM culture medium and infected at an MOI of 10 for 3 to 6h. At the end of the infection, cell supernatant was removed, and cells were fixed with a 4% PFA solution for 10 min at 37 °C. When specified, plasma membrane was stained with Wheat Germ Agglutinin-Alexa Fluor 647 (WGA) (ThermoFisher, W32466) at 1:250 dilution in HBSS and incubated for 10 min in the dark. Permeabilization was performed by incubating cells for 10 min in PBS containing 0.1% Triton X-100. To block unspecific binding of antibodies, cells were incubated in PBS-T (PBS + 0.1% Tween 20), containing 2% BSA, for 30 min. Cells were incubated with primary antibodies (1:100) anti-NE (abcam, ab314916), anti-MPO (abcam, ab208670), anti-Ly6G (Biolegend, 127602), overnight at 4 °C in PBS-T + 2% BSA solution and detected with the appropriate secondary antibodies (1:1000) Alexa Fluor 555-conjugated donkey anti-rabbit (ThermoFisher, A31572) or AlexFluor 647-conjugated goat anti-rat (ThermoFisher, A21247), for 1 hour at room temperature. DNA was counterstained with DAPI (1:1000). Stained cells were imaged using Zeiss AxioVision Z1 microscope and analysed with Zen 2.3 Blue Version (Carl Zeiss MicroImaging GmbH, Germany).

### Quantification of NETs

Neutrophils were plated at a density of 5.10^5^ cells on glass bottom Ibidi chambered coverslips and infected at an MOI of 10 for 3 to 6h in Opti-MEM culture medium supplemented with SYTOX-Green dye (100ng/mL). At the end of the experiment, DAPI was added to the medium and NETs were visualised in at least six random images (20x magnification) per sample. NETs were quantified with ImageJ software by counting the number of SYTOX-positive cells and the associated DNA area for each SYTOX-positive cells. The percentage of SYTOX-positive cells was obtained by dividing the SYTOX-positive counts to the total number of cells.

### Kwik-Diff staining of BALs

BALs were centrifuge at 300xg for 10min, and pellet was resuspended in 1X PBS supplemented with 3mM EDTA. Absolute cell number was determined with automated countess 3 cell counter (Invitrogen) with trypan blue cell death exclusion method and cell density was adjusted at 2.10^6^ / mL by adding 1X PBS + 3mM EDTA. Cells (100µL) were plated in coated slides (Epredia, 12026689) by centrifugation at 500xg for 5min using a Cytospin (ThermoFisher, Shandon), before staining with Kwik-Diff (Epredia Shandon, 10435310), according to the manufacturer’s instructions. Briefly, slides were fixed in Kwik-Diff reagent 1 (5 seconds) and stained sequentially by incubation into Kwick-Diff reagent 2 (1 min) and reagent 3 (30 seconds). Excessive staining was removed by performing several gentle washes (distilled water) and air dried for 30min at room temperature. Slides were mounted using hardset vectashield and imaged with AxioVision Z1 microscope.

### Scanning and Transmission Electron Microscopy experiments

For Scanning Electron Microscopy observations, cells were fixed with 2.5% glutaraldehyde in 0.2M cacodylate buffer (pH 7.4). Preparations were then washed three times for 5min in 0.2M cacodylate buffer (pH 7.4) and washed with distilled water. Samples were dehydrated through a graded series (25 to 100%) of ethanol, transferred in acetone and subjected to critical point drying with CO2 in a Leica EM CPD300. Dried specimens were sputter-coated with 3 nm platinum with a Leica EM MED020 evaporator and were examined and photographed with a FEI Quanta FEG250.

For Transmission Electron Microscopy, samples were fixed using a final concentration of 2.5% glutaraldehyde and 2% paraformaldehyde in 0.1 M Sorensen buffer (pH 7.2; EMS, EMS11600-05), added to the culture medium in a volume-to-volume ratio for 15 minutes at room temperature. Following this primary fixation, the cells underwent secondary fixation with the same solution (2.5% glutaraldehyde, 2% paraformaldehyde in 0.1 M Sorensen buffer, pH 7.2) for 2 hours at room temperature and were then post-fixed at room temperature with 1% osmium tetroxide (OsO₄) and 1.5% potassium ferricyanide K3Fe(CN)6 in 0.1 M cacodylate buffer with 2 mM CaCl₂. Next, cells were treated overnight at 4°C with 2% aqueous uranyl acetate, dehydrated through a graded ethanol series, and embedded in Epon resin. After 48 hours of polymerization at 60°C, ultrathin sections (80 nm) were mounted onto 200-mesh Formvar-carbon-coated copper grids. Sections were then stained with Uranyless and lead citrate.Grids were examined using a transmission electron microscope (TEM; Jeol JEM-1400, JEOL Inc., Peabody, MA, USA) operated at 80 kV, and images were acquired with a digital camera (Gatan Orius, Gatan Inc., Pleasanton, CA, USA).

### Statistical analysis

All data are from at least three independent experiments. Statistical analyses were performed with Prism (GraphPad Software, v10.4.0). All statistical tests and p-values are mentioned in the figure legends.

